# Perception of Rigidity in Three- and Four-Dimensional Spaces

**DOI:** 10.1101/2022.11.09.515874

**Authors:** Dongcheng He, Dat-Thanh Nguyen, Haluk Ogmen, Shigeaki Nishina, Arash Yazdanbakhsh

**Affiliations:** Department of Electrical & Computer Engineering, Laboratory of Perceptual & Cognitive Dynamics, University of Denver, Denver, CO, USA; Department of Psychological and Brain Sciences, Computational Neuroscience and Vision Lab, Center for Systems Neuroscience, Boston University, Boston, MA, USA; Honda Research Institute Japan Co., Ltd., Wako, Japan

**Author notes:** Equal contribution (Dongcheng He, Dat-Thanh Nguyen).

## Abstract

Our brain employs mechanisms to adapt to changing visual conditions. In addition to natural changes in our physiology and those in the environment, our brain is also capable of adapting to “unnatural” changes, such as inverted visual-inputs generated by inverting prisms. In this study, we examined the brain’s capability to adapt to hyperspaces. We generated four spatial-dimensional stimuli in virtual reality and tested the ability to distinguish between rigid and non-rigid motion. We found that observers are able to differentiate rigid and non-rigid motion of tesseracts (4D) with a performance comparable to that obtained using cubes (3D). Moreover, observers’ performance improved when they were provided with more immersive 3D experience but remained robust against increasing shape variations. At this juncture, we characterize our findings as “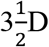 perception” since, while we show the ability to extract and use 4D information, we do not have yet evidence of a complete phenomenal 4D experience.

## 1. Introduction

Visual information about our environment is projected as two-dimensional images on our retinae. By using various cues, such as disparity, motion parallax, perspective, and shading, the visual system *constructs* three-dimensional percepts from these two-dimensional projections. Hence, our brains are capable of augmenting the dimensionality of their inputs.

A question arises as to whether our brains are limited to three-dimensional representations. On the one hand, one may argue that, since we live in a threedimensional environment, our brains are evolved to represent three-dimensional percepts only. On the other hand, our brains show *plasticity*, i.e., the capability of adapting to changing conditions. Vision does not start in a mature state in infants; rather it is through visual adaptation and developmental processes that it reaches its mature state. Visual development requires interactions with the environment. The plasticity associated with these interactions is not limited to infants but also continues in adulthood. For example, using inverting lenses, Stratton (1896) showed that we can adapt to image inversion and carry out complex tasks successfully, such as riding a bicycle in city streets. Given this ability to adapt to “unnatural conditions”, one may postulate that, with sufficient experience with higher-dimensional (four or more spatial dimensions) stimuli, our brains may be able to adapt to these higher spatial-dimensional stimuli. This hypothesis can be tested experimentally. Even though our environment is three-dimensional, mathematically we can generate higher-dimensional stimuli (“hyperspaces”) and expose subjects to these stimuli.

In fact, previous studies have shown that human subjects can perform better than chance in hyperspace reasoning or hyperdimensional object recognition tasks (review: Ogmen et al., 2020). In Aflalo and Graziano (2008)’s study, subjects were found to effectively learn to navigate in 4D mazes constructed from 2D displays. Ambinder et al. (2009) found that subjects can build geometrical concepts of angle and distance when exposed to 4D tetrahedrons. Wang (2014 a; 2014b) investigated if subjects were able to measure the hypervolume of 4D objects by learning rotated 4D objects and found that subjects showed the ability to judge the actual hypervolume. Miwa et al. (2018) in their study presented different perspectives of the 3D projections of the 4D objects and let the subjects actively interact with the presentation by manually controlling the rotation of objects via a Virtual Reality (VR) system. Performance suggested that subjects were able to reconstruct the 4D space based on multiple 3D projections. Most of the previous studies investigated hyperspace perception by designing stimuli and tasks related to spatial reasoning, spatial relations, and perspective learning. Motion is a key source of information in visual perception and motion parallax plays a major role in perceiving the third dimension from twodimensional stimuli. Whether and how motion can be used to generate higher-dimensional percepts remains largely unexplored. We are often able to identify stable objects across transformations and have strong subjective impressions of the transformations themselves. This suggests that the brain is equipped with sophisticated mechanisms for inferring both object constancy and objects’ causal history. The ability to perceive and classify the rigidity of object transformation is thus an important aspect of spatial representations.

In this study, we used a VR system to present subjects with hypercubes undergoing either rigid or non-rigid motion and tested their ability to judge the rigidity of the stimuli.

## 2. Materials and Methods

### 2.1. Subjects

Six students (two women and four men; age: M[SD]=21[3.16] years), including one of the authors, from the University of Denver participated in the first experiment and all participants had a normal or corrected-to-normal vision. Five subjects (one woman and four men; age: M[SD]=22.3[2.62] years) who attended experiment 1 participated in the second experiment. These experiments followed a protocol approved by the University of Denver Institutional Review Board for the Protection of Human Subjects. Each observer gave written consent before the experiment.

### 2.2. Apparatus

In this experiment, we presented our stimulus using an HTC VIVE VR headset released in 2017. The VR headset possesses two rectangular screens with circular lenses for the two eyes, each with a diagonal size of 91.4 mm. The resolution of each screen was 1200×1080 with a 90 Hz refresh rate. The approximate pupil-to-lens distance was 18 mm. The display of the device is controlled by the development environment Unity3D.

### 2.3. Description of Stimuli

The stimuli were distorted 3D cube and 4D cube wireframe objects (Figure 1). A 3D object consisted of 8 vertices and 12 edges, and a 4D object consisted of 16 vertices and 32 edges. The objects were presented in a 3D virtual space using the VR headset. For the 4D object, we presented its 3D projection. We made the distorted object shape by shifting the position of each vertex of a regular cube with side lengths of 100 cm in a random direction by one of the three amounts: 12, 18, or 24 cm, representing the three irregularity levels, as shown in Figure 2.

**Figure 1:**
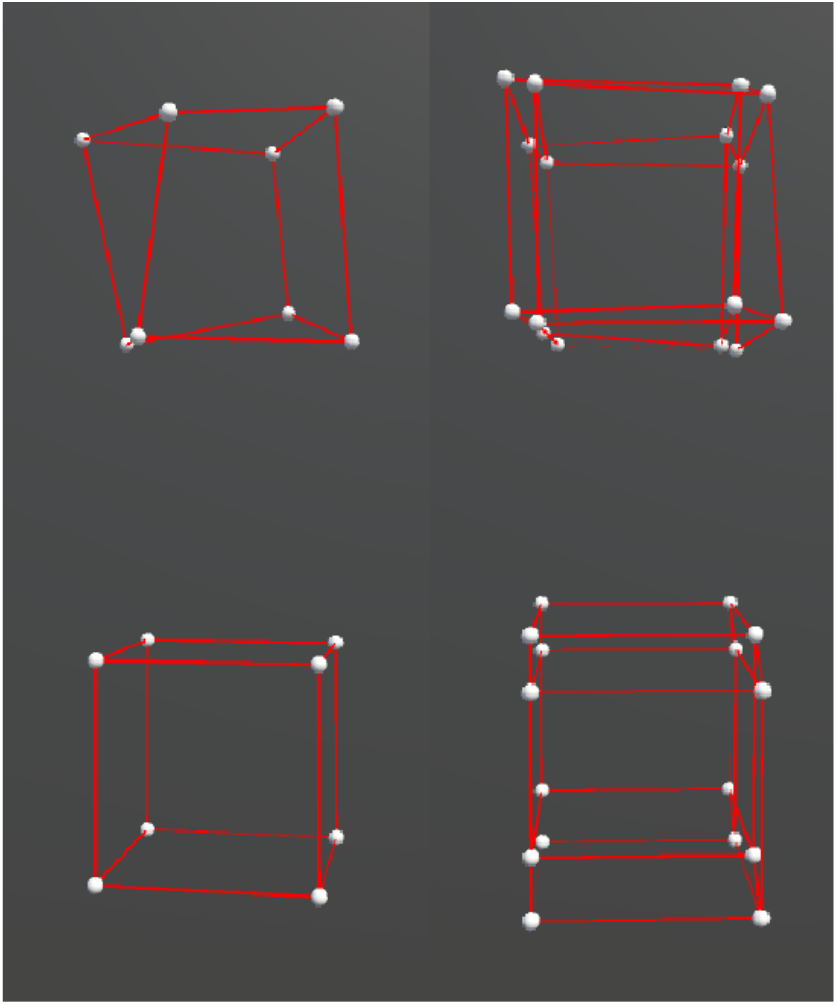
Distorted cubes used in the experiment. The left and right columns show the cubes in 3D and 4D respectively. The bottom row contains the original cubes, and the top row contains the cubes after adding the distortion.

**Figure 2:**
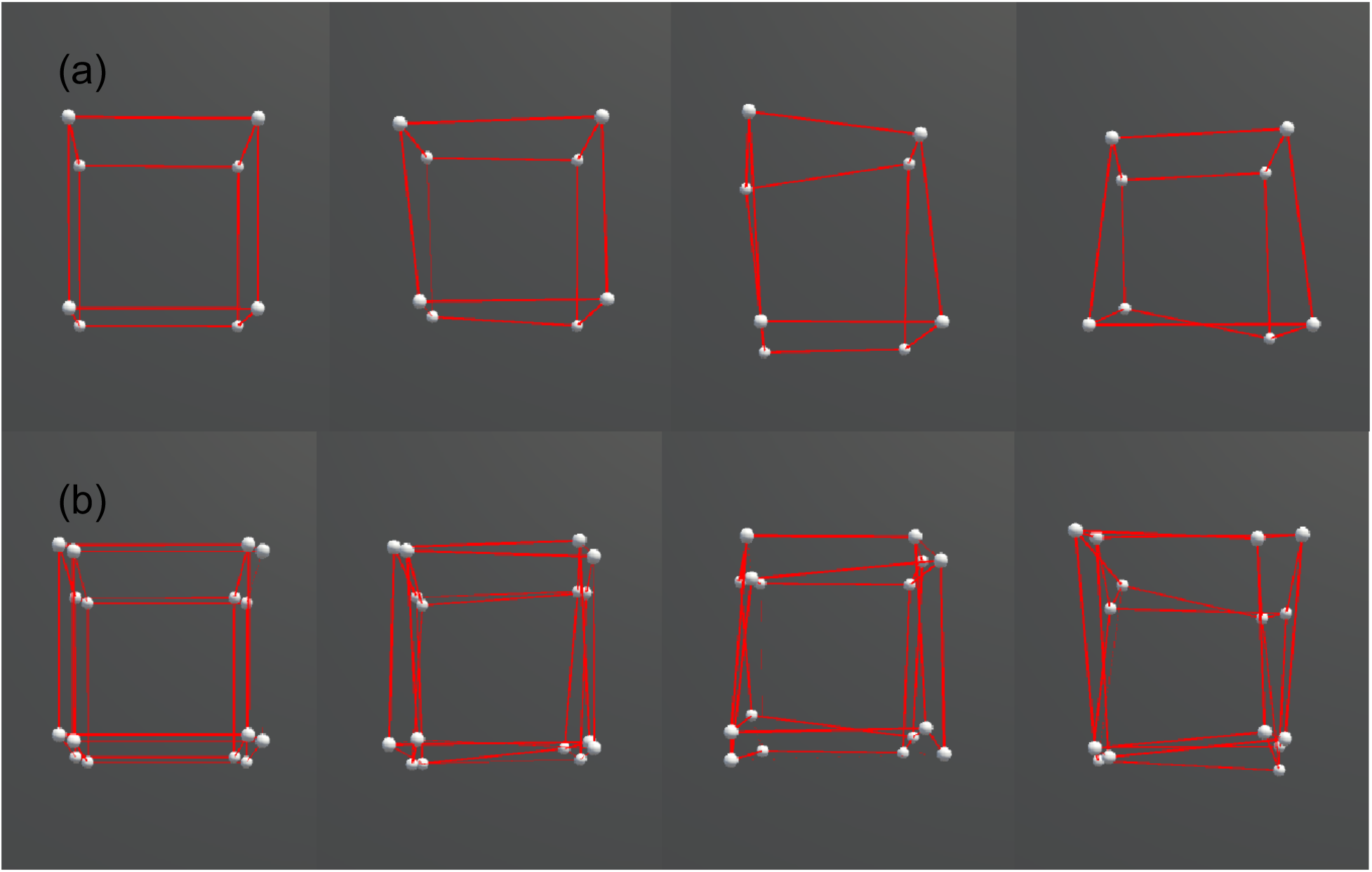
Comparison between different ranges of irregularity. From left to right: 0, 0.12, 0.18, 0.24.**(a)** 3D cubes. **(b)** 4D cubes.

We presented two types of object motion in the experiment: rigid and non-rigid. For both two types, a slight random motion was applied to each vertex to enhance the stereoscopy of the object. Specifically for the rigid motion, the entire object rotated repeatedly over a fixed angular range around the x and y axes, and the axial information is depicted in Figure 3. Generalized to the 4D cubes, it was a plane, instead of a line, that serves as the rotation axis, and thus the 4D cubes rotated along y-z, x-z, and x-y planes. Whereas for the non-rigid motion, in addition to the same rotation as the rigid motion, non-rigidity was made by simultaneously deforming the object along random directions and adding the axial displacements via shearing to the x, y, or z axis, as shown in Figure 4. The mathematical descriptions of the stimuli can be found in the Appendix I.

**Figure 3:**
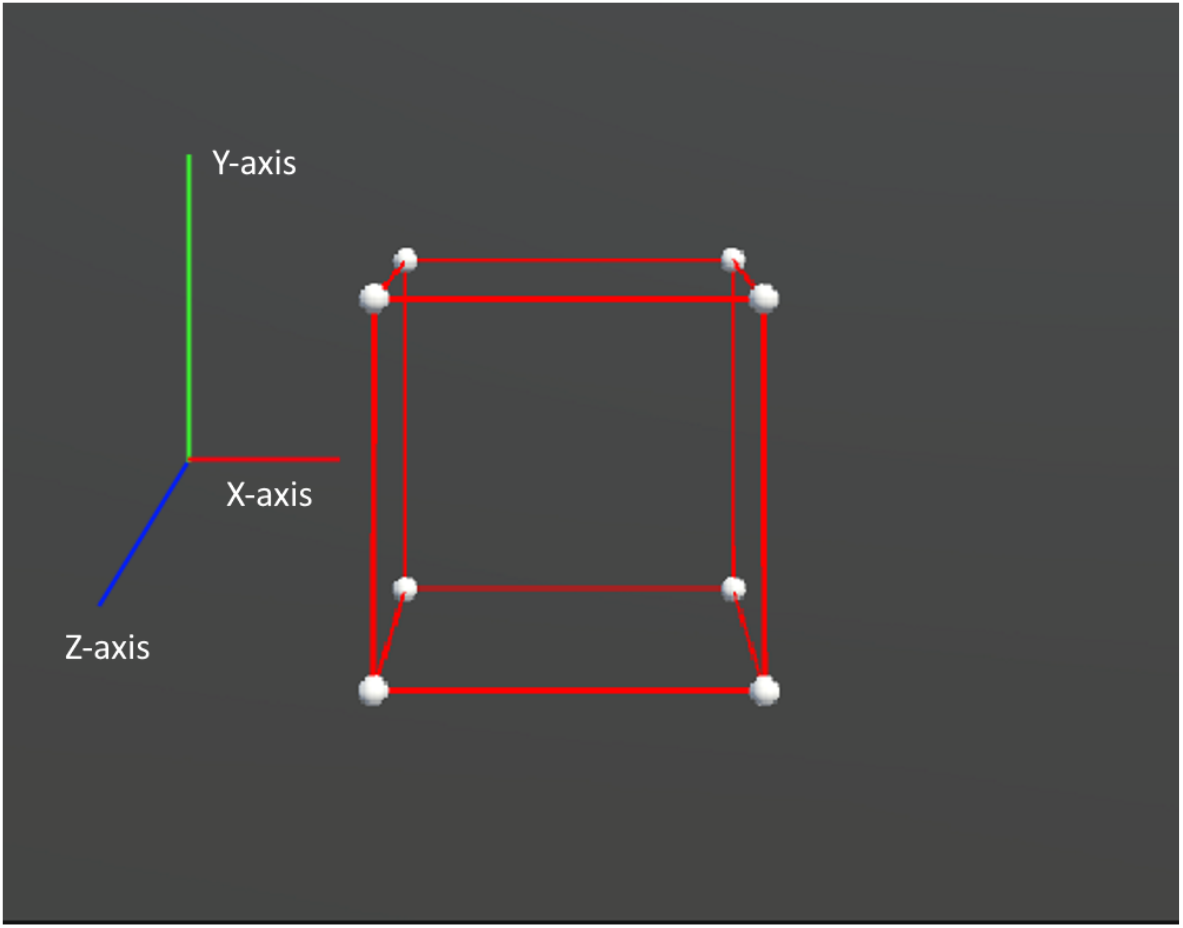
Axis legend for the virtual experiments. The X axis represents left to right, the Y axis represents up to down, and the Z axis represents in to out.

**Figure 4:**
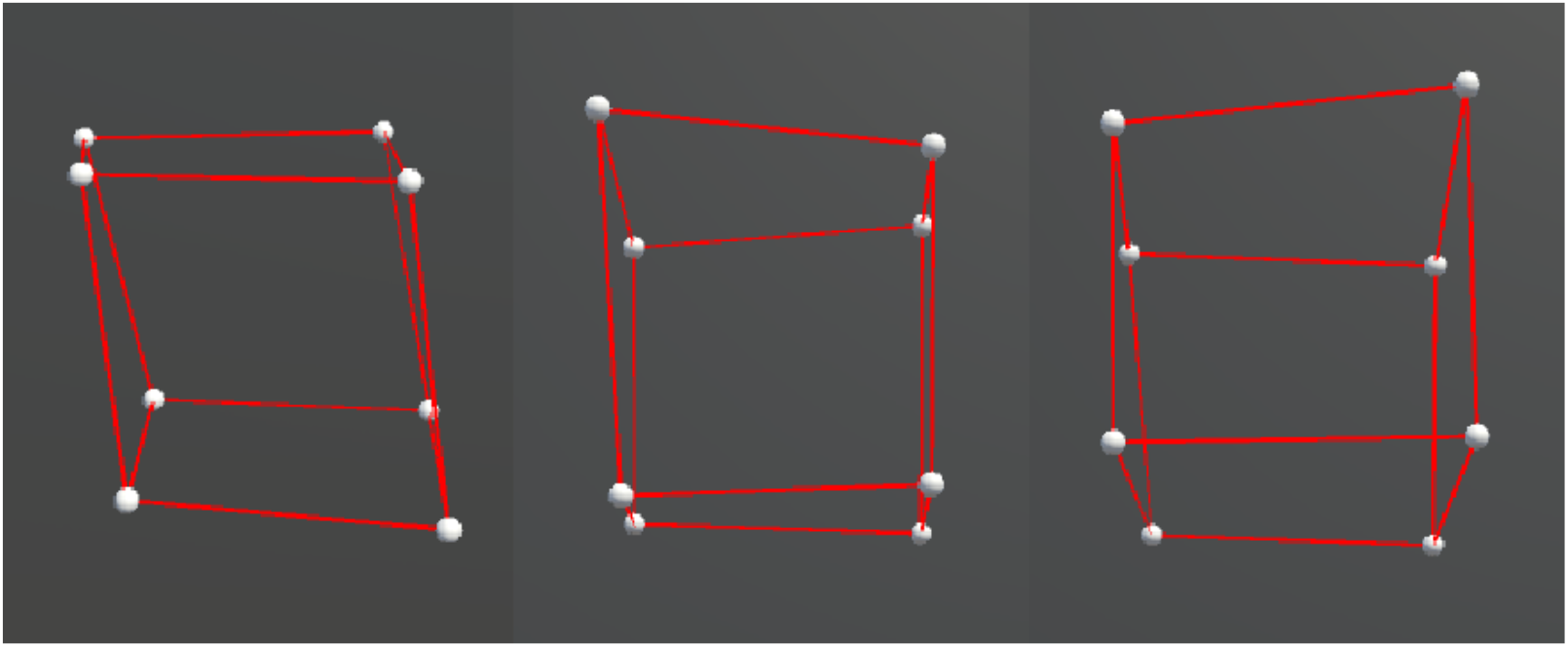
Examples of each displacement axis condition, from left to right: x-axis (left-right) displacement, z-axis (in-out) displacement, and y-axis (up-down) displacement.

In Experiment 1, The distance from the subject’s eyes to the center of the object in the virtual space was fixed. The objects were presented in perspective projection. In addition to the perspective cue, depth information in the 3D presentation was collaboratively given by (1) a binocular cue provided by the stereoscopic effect of VR and (2) a structure-from-motion cue generated by randomly rotating the entire object. Motion parallax cue was also available in Experiment 2, in which the subjects were allowed to move their heads actively.

### 2.4. Experimental Design

#### 2.4.1. Experiment 1

In a normally illuminated room, subjects sat on a chair and observed the stimuli via the VR headset that was placed on a rack fixed on the table. The viewpoint of the observation was fixed toward the front view of the stimulus regardless of the headset’s position and rotation. As shown in Figure 5, beginning of each trial, two cubes were presented in a top-bottom manner, in which one had rigid motion and the other had non-rigid motion. With text notification, subjects were guided to determine which one of the two cubes has rigid motion by pressing one of the two buttons of a mouse (press the left button to report the top and the right button to report the bottom cube). This window would last for up to 3 seconds. Timing-out would trigger the next trial automatically and thus overtime responses would not be recorded. Once reported, the next window would come when subjects were asked to report their confidence in the previous decision by pressing a key from 1 to 5 on a keyboard, where 1 means purely guessing and 5 means firm belief. The next trial would come as soon as they pressed a key. A video demo showing a trial can be found on the following link: https://github.com/hedch/4D_reconstruction. A session consisted of objects of two dimensionalities (3D and 4D) in motion along three displacement axes (x, y, and z), and by three irregularity levels (0.12, 0.18, and 0.24). Each case was repeated by three times, and thus there were 54 trials in a session. 3D and 4D trails were separated by text instruction and together with a time interval of 5 seconds. Within each dimensional condition, the sequence of presentations was randomized in a counter-balanced order. Each subject took three sessions of experiments, in which the first session was for training purposes and thus wasn’t included in the analysis.

**Figure 5:**
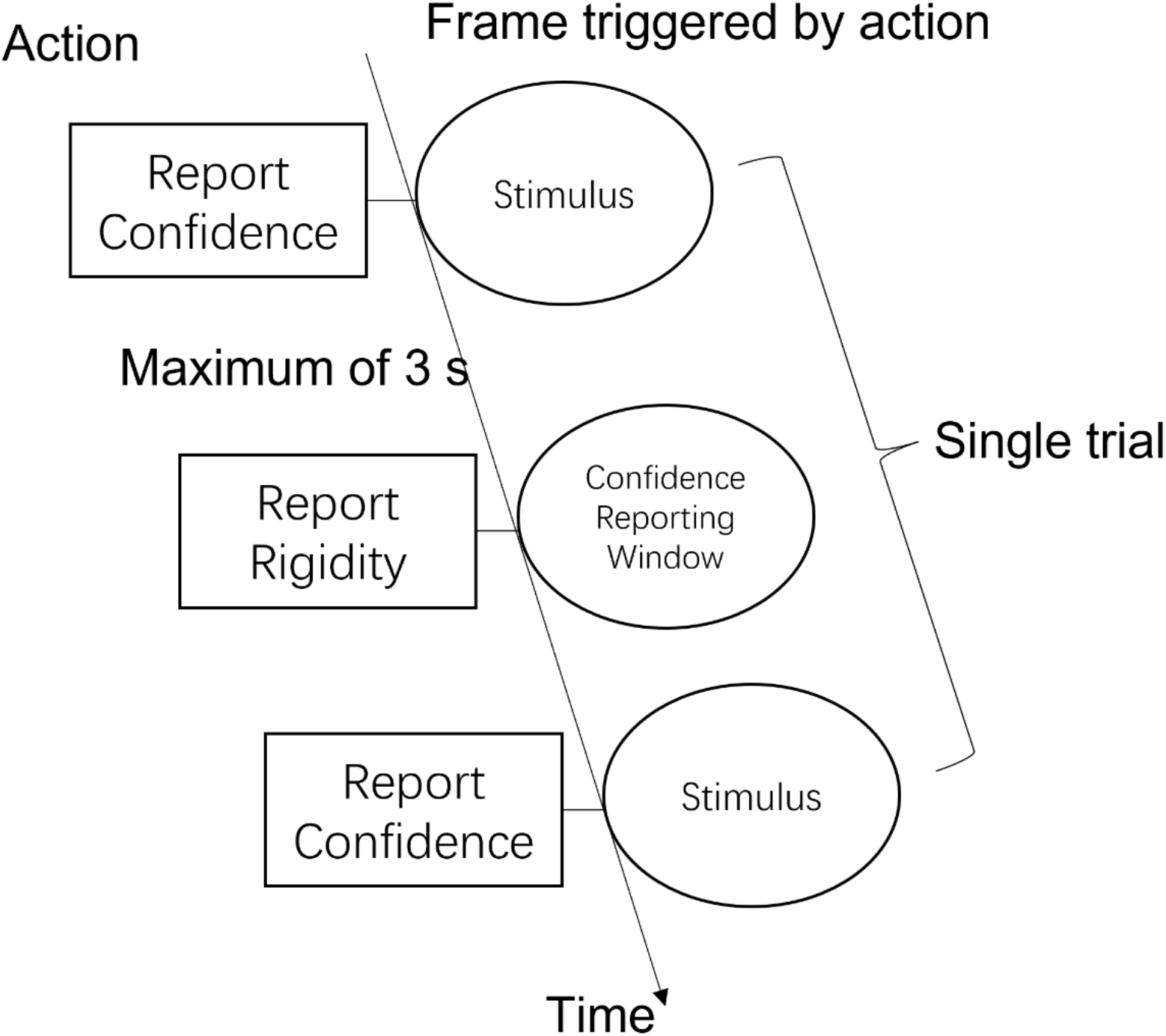
Schematic of a trial.

#### 2.4.2. Experiment 2

The single trial procedure was same as the experiment 1. However, in this experiment, the x-axis displacement was removed. Therefore, a session consisted of objects of two dimensionalities (3D and 4D) in motion along two displacement axes (y and z), and by three irregularity levels (0.12, 0.18, and 0.24). Each case was repeated by seven times, and thus there were 84 trials in a session. Due to the previous experience of all subjects, no training session was done, and each subject was asked to finish two sessions of experiments with about ten minutes of resting between them. Importantly, observers wore the headset during the experiment and their viewpoint in the VR environment was changed accordingly as they move or rotate their heads. All other experimental settings were the same as Experiment 1.

### 2.5. Results

#### 2.5.1. Experiment 1

Among the six subjects, one had a performance lower than 60% correct in the experiments and whose data were therefore excluded from the analysis. The percentages of the correctness of the other five subjects are 77.78%, 65.74%, 79.63%, 75.93%, and 64.81% respectively.

Figure 6 shows accuracy in terms of percent-correct responses for different stimulus dimensions and displacement axes. For the fixed-headset condition, a repeated-measures ANOVA with displacement axes and dimensions as main factors showed a significant effect of displacement axis [F(2,8)=34.59, p<0.01] on accuracy. Specifically, as detailed in Table 1, the performance under the x-displacement condition was remarkably better than the other two displacement axes. The effect of dimension was found to be non-significant on accuracy [F(1,4)=0.25, p=0.64]. There was no significant interaction between displacement axes and dimension [F(2,8)=2.48, p=0.14].

**Figure 6:**
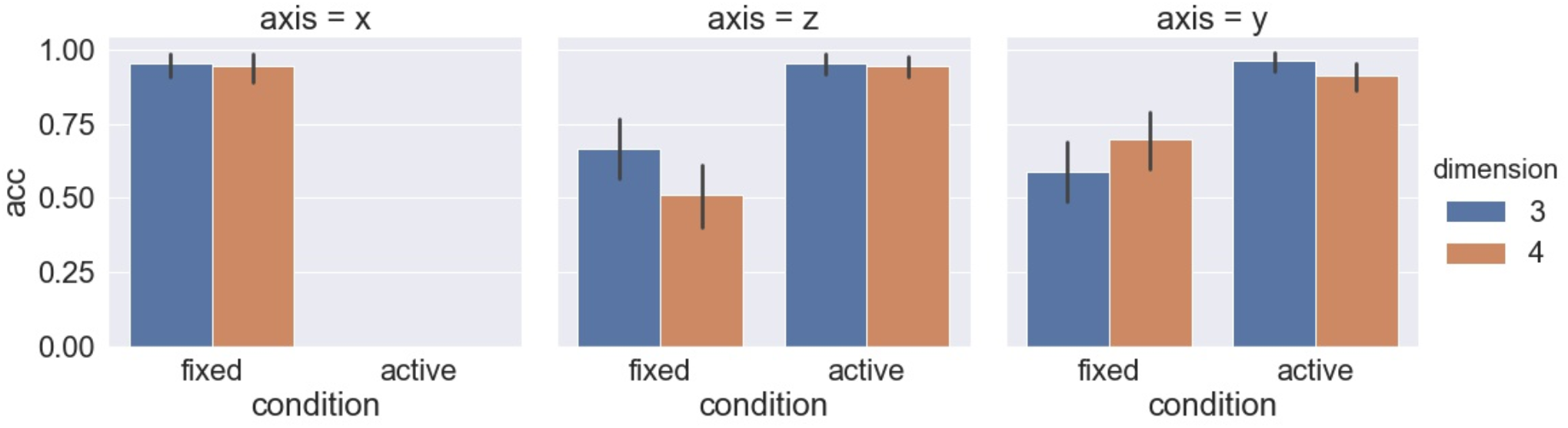
Percent correct in each displacement axis condition.

**Table 1:**
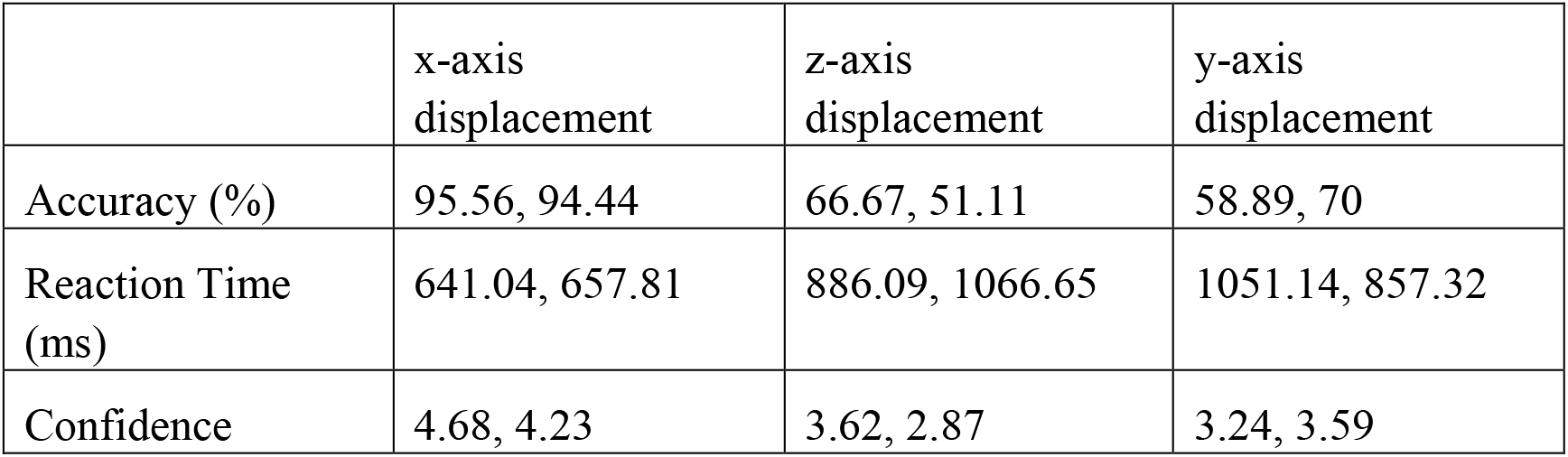
Fixed Headset Condition: 3D measurement, 4D measurement in means.

Figure 7 shows the reaction time by the dimensions and displacement axes. For the fixed-headset condition, a repeated-measures ANOVA showed a significant effect of displacement axes [F(2,8)=14.36, p<0.01] on reaction time. Coinciding with the accuracy, a shorter time was spent in the rigidity discrimination task when the vertexes were displaced along the x-axis compared to the other two axes, as shown in Table 1. The effect of dimension was found to be non-significant on reaction time [F(1,4)=0.01, p=0.91]. However, the interaction between the displacement axis and dimension was found to be significant [F(2,8)=14.67, p<0.01], which can be interpreted by the inverse effects of dimension under the y-axis and z-axis displacements.

**Figure 7:**
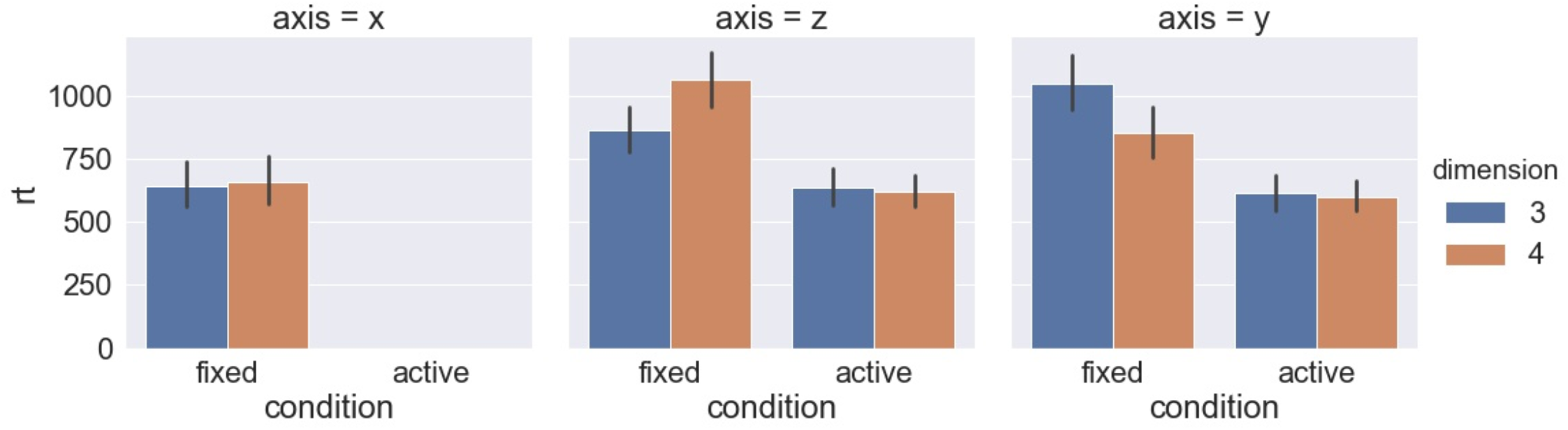
Reaction time (ms) in each displacement axis condition.

Subjects’ reported confidence levels are shown in Figure 8, and a repeated-measures ANOVA showed a significant effect of the displacement axis [F(2,8)=19.09, p<0.01] on the confidence level. In general, subjects were more confident when the cubes m along the x-axis in contrast to the other two axes, which was consistent with their behaviors reflected by the reaction time and accuracy. Also, dimension was found to have no significant effect on confidence level [F(1,4)=3.39, p=0.14], and no significant interaction between displacement axis and dimension was detected [F(2,8)=2.62, p=0.13].

**Figure 8:**
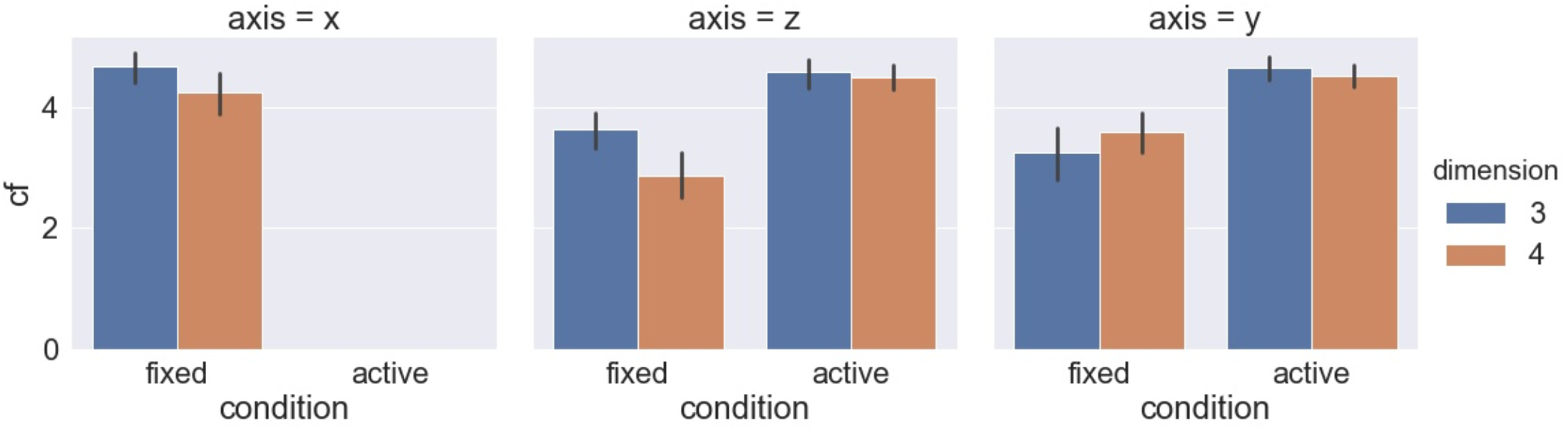
Confidence level reported in each displacement axis condition.

Measurements with respect to irregularities are shown in Figures 9–11. Repeated-measures ANOVAs with irregularity levels, dimensions, and headset conditions as main factors showed that irregularity level has no significant effects on accuracy [F(2,8)=0.23, p=0.8], reaction time [F(2,8)=0.54, p=0.6], and confidence [F(2,8)=1.69, p=0.24]. No interaction between irregularity and other factors was found to be significant in all tests.

**Figure 9:**
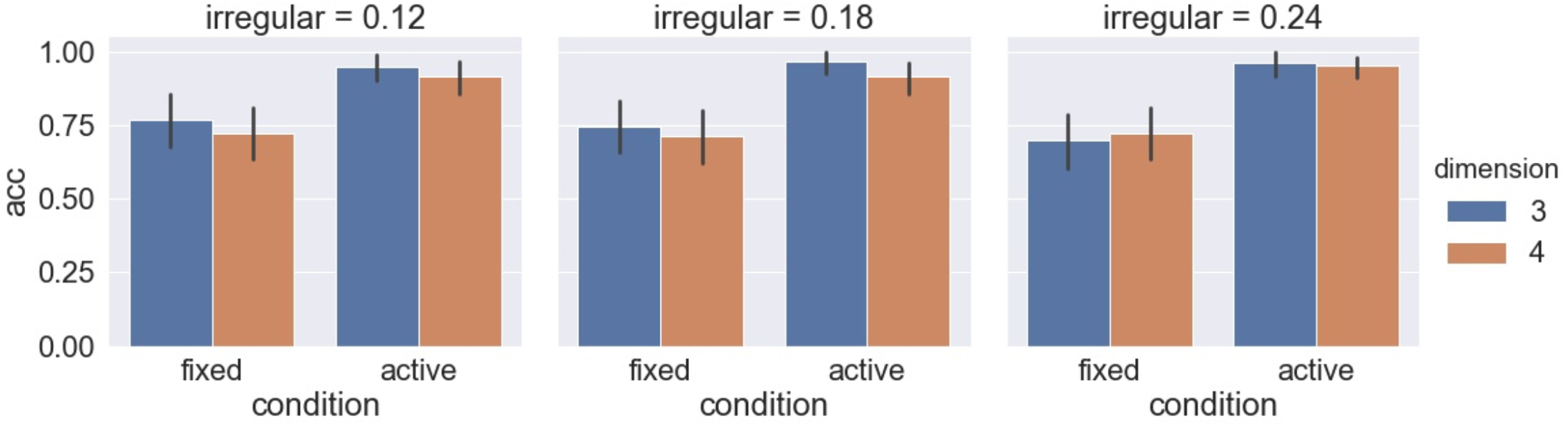
Percent correct in each irregular condition.

**Figure 10:**
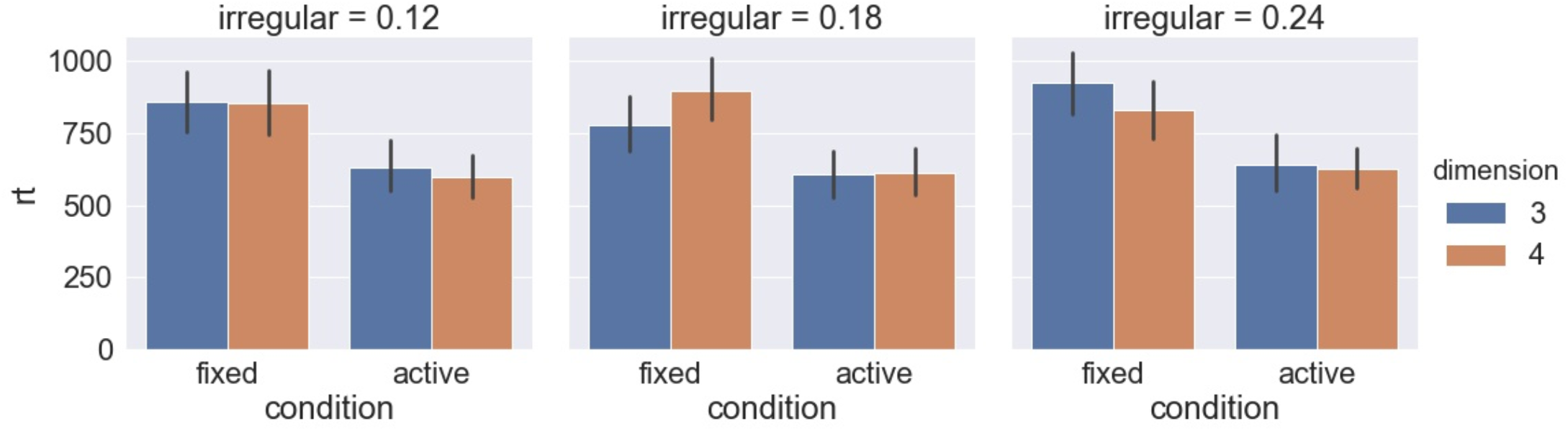
Reaction time (ms) in each irregular condition.

**Figure 11:**
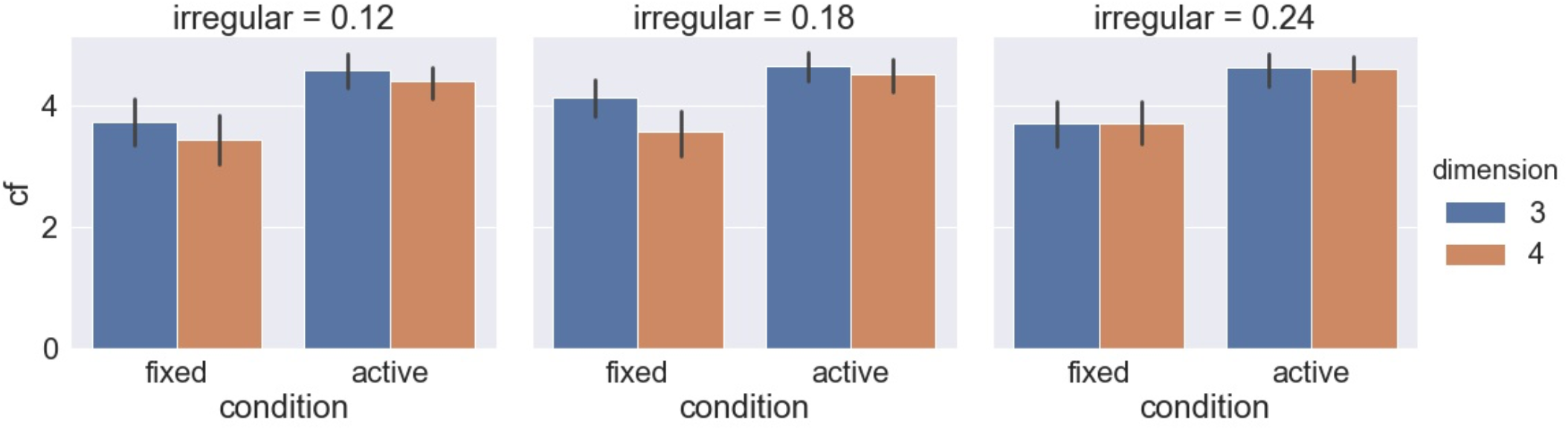
Confidence level reported in each irregular condition.

#### 2.5.2. Experiment 2

##### Reported Measurements

The five subjects who participated in Experiment 2 were identical to the five subjects who gave an acceptable performance in Experiment 1, and their percent correct performance herein were 100%, 91.67%, 99.07%, 91.67%, and 89.81% respectively.

Figure 6 shows the accuracy of the dimensions and displacement axis. For the active headset condition, a repeated-measures ANOVA with displacement axes and dimensions as main factors was conducted but no effect was found to be significant on accuracy [Axis: F(1,4)=1.81, p=0.25; Dimension: F(1,4)=1.17, p=0.34; Axis×Dimension: F(1,4)=1.75, p=0.26]. Figure 7 shows the reaction time by the dimensions and displacement axes. Similarly, a repeated-measures ANOVA detected no significant effect from the displacement axis and the dimension on reaction time [Axis: F(1,4)=0.14, p=0.73; Dimension: F(1,4)=1.09, p=0.35; Axis×Dimension: F(1,4)=0.03, p=0.88]. Moreover, as shown in Figure 8, unsurprisingly given the results in accuracy and reaction time, the same ANOVA didn’t detect any significant effect on confidence level neither [Axis: F(1,4)=0.05, p=0.83; Dimension: F(1,4)=0.18, p=0.69; Axis×Dimension: F(1,4)=0.67, p=0.46]. Detailed measurements can be found in Table 2.

**Table 2:** Active Headset Condition: 3D measurement, 4D measurement in means.

|  | z-axis displacement | y-axis displacement |
| --- | --- | --- |
| Accuracy (%) | 95.49, 94.61 | 96.35, 91.43 |
| Reaction Time (ms) | 637.81, 622.3 | 615.14, 601.89 |
| Confidence | 4.57, 4.49 | 4.66, 4.52 |

Measurements with respect to irregularities are shown in Figures 9–11. Repeated-measures ANOVAs with irregularity levels, dimensions, and headset conditions as main factors showed that irregularity level had no significant effects on accuracy [F(2,8)=1.33, p=0.32], reaction time [F(2,8)=0.25, p=0.78], and confidence [F(2,8)=0.44, p=0.66]. No interaction between irregularity and other factors was found to be significant in all tests.

##### Comparisons between Fixed and Active Headset Conditions

As explained, preliminary data showed that active exploration would improve the performance in this rigidity motion discrimination task. Considering the ceiling accuracy observed in Experiment 1 under the x-axis displacement condition, in Experiment 2 we tested under the y-axis and z-axis conditions only. Therefore, to test the improvement in performance raised by the active headset statistically, here we first tested the effect of observation conditions (fixed headset and active headset) under the y-axis and z-axis conditions only. Repeated-measures ANOVAs with observation conditions, displacement axis, and dimensions as main factors found that the observation condition had significant effects on all the three reported measurements [Accuracy: F(1,4)=123.74, p<0.01; Reaction time: F(1,4)=15.17, p=0.02; Confidence level: F(1,4)=34.67, p<0.01]. Referring to the values shown in Tables 1 and 2, significant improvements took place in accuracy, reaction time, and confidence. In addition, compensated by these improvements, these performances can be comparable to how subjects behaved in Experiment 1 under the x-axis condition, as another repeated-measures ANOVA including the data groups in Experiment 2 involving y- and z-axis and the data group in Experiment 1 involving x-axis only detected no significant difference among the three data groups [Accuracy: F(1,4)=1.18, p=0.25; Reaction time: F(1,4)=0.14, p=0.73; Confidence level: F(1,4)=0.05, p=0.83].

##### Head Movements during Subjects’ Active Explorations

To analyze the correlation between the head movement and the performance, we recorded the visual angles along the x, y, and z axes calculated by the angles between the headset’s viewing direction and the vector from the headset’s position to the cube’s position. In each trial, angles were scanned by approximately 18 Hz, and therefore about 54 data points were recorded during the 3-second observation period in each trial. We then calculated the median and the percentile range of angles collected along each axis in each trial to investigate the effect of viewpoint and its change on performance. The percentile ranges were calculated by the third quarter value subtracted from the first quarter value. We didn’t use the maximum and minimum as they are sensitive to outliers. As shown in Figures 12–14, we found that, for all subjects, the ranges of viewpoints within trials were mostly small and they were narrow across all trials. Besides, we didn’t find effective correlations between the range of viewpoints and the performance, as detailed in Table 3.

**Figure 12:**
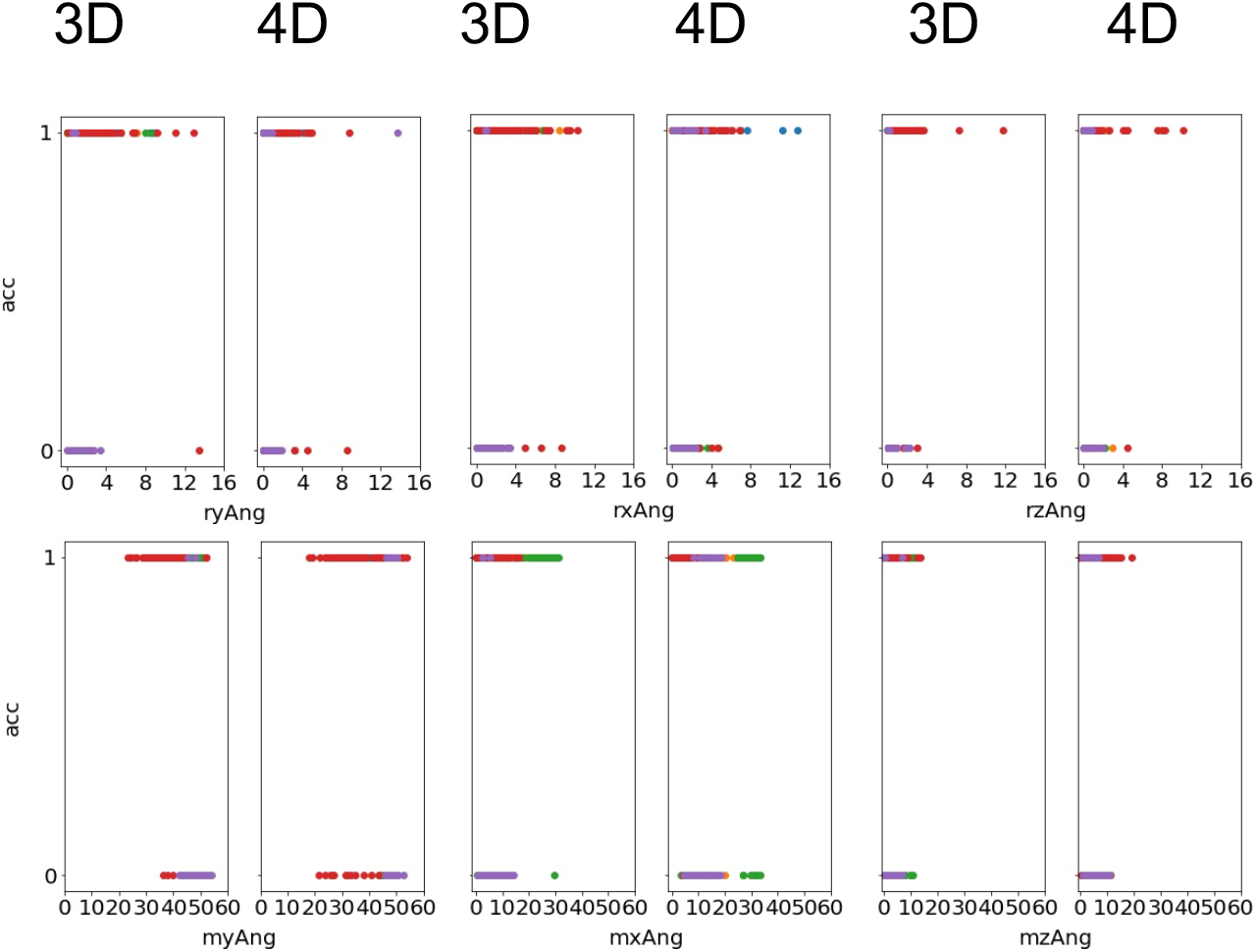
The correlation between the accuracy (1/0 represents correct/false) and the angles (deg). Each dot represents a statistical summary of a trial. Each color correponds to a subject. The horizontal axes labels indicate the shearing axis (x, y, or z), and the statistics (range or median). For example, “ryAng” indicates the range of headset’s rotational angles along the y axis during a trial, which was calculated by the third quarter value minus the first quarter value. “mxAng” indicates the median of angles between the headset positions and the cube relative to the x axis during a trial. One outlier data point with rzAng=27.25° was removed from Figure 12 to 14.

**Figure 13:**
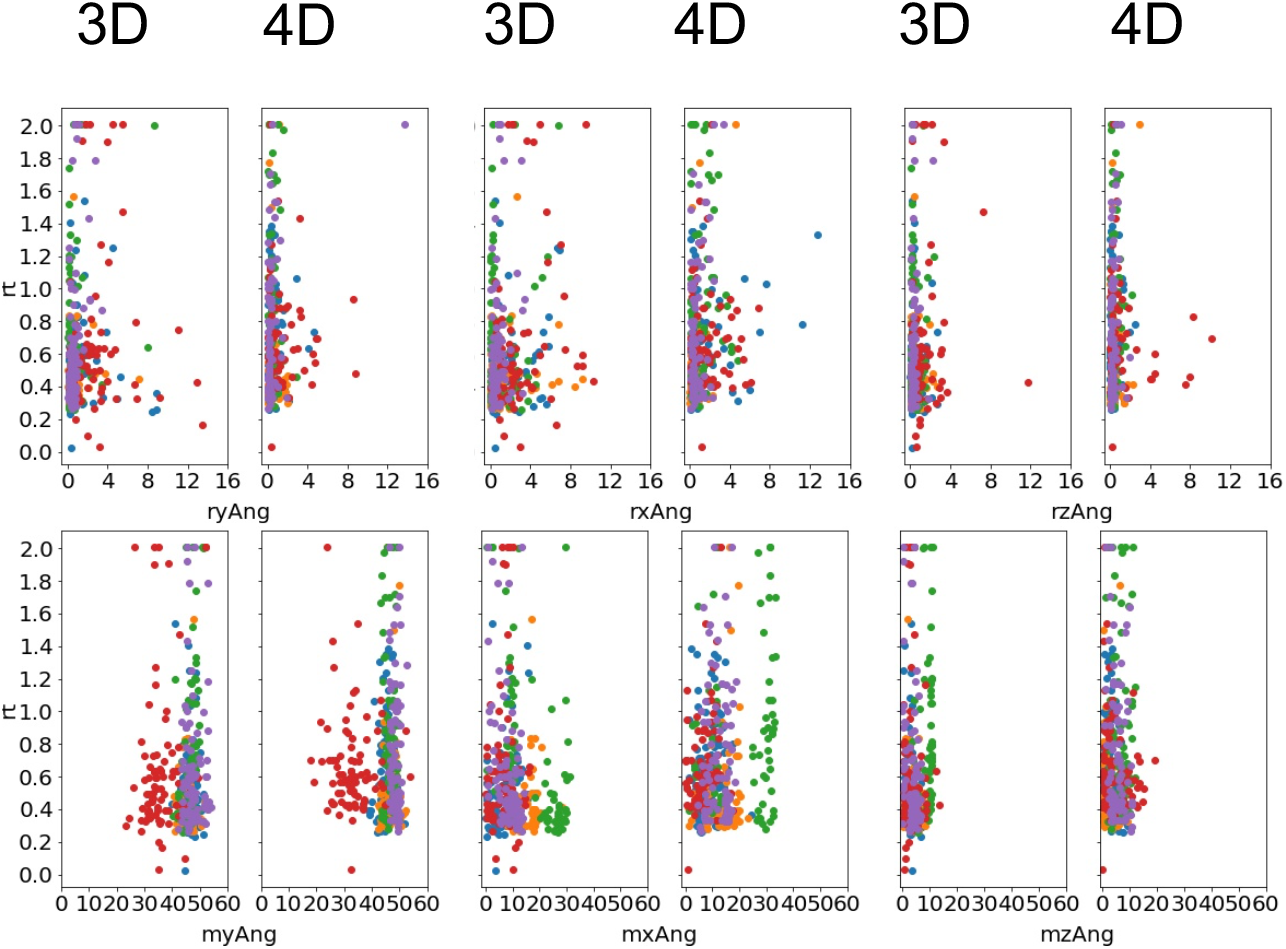
The correlation between the reaction times (s) and the angles (deg).

**Figure 14:**
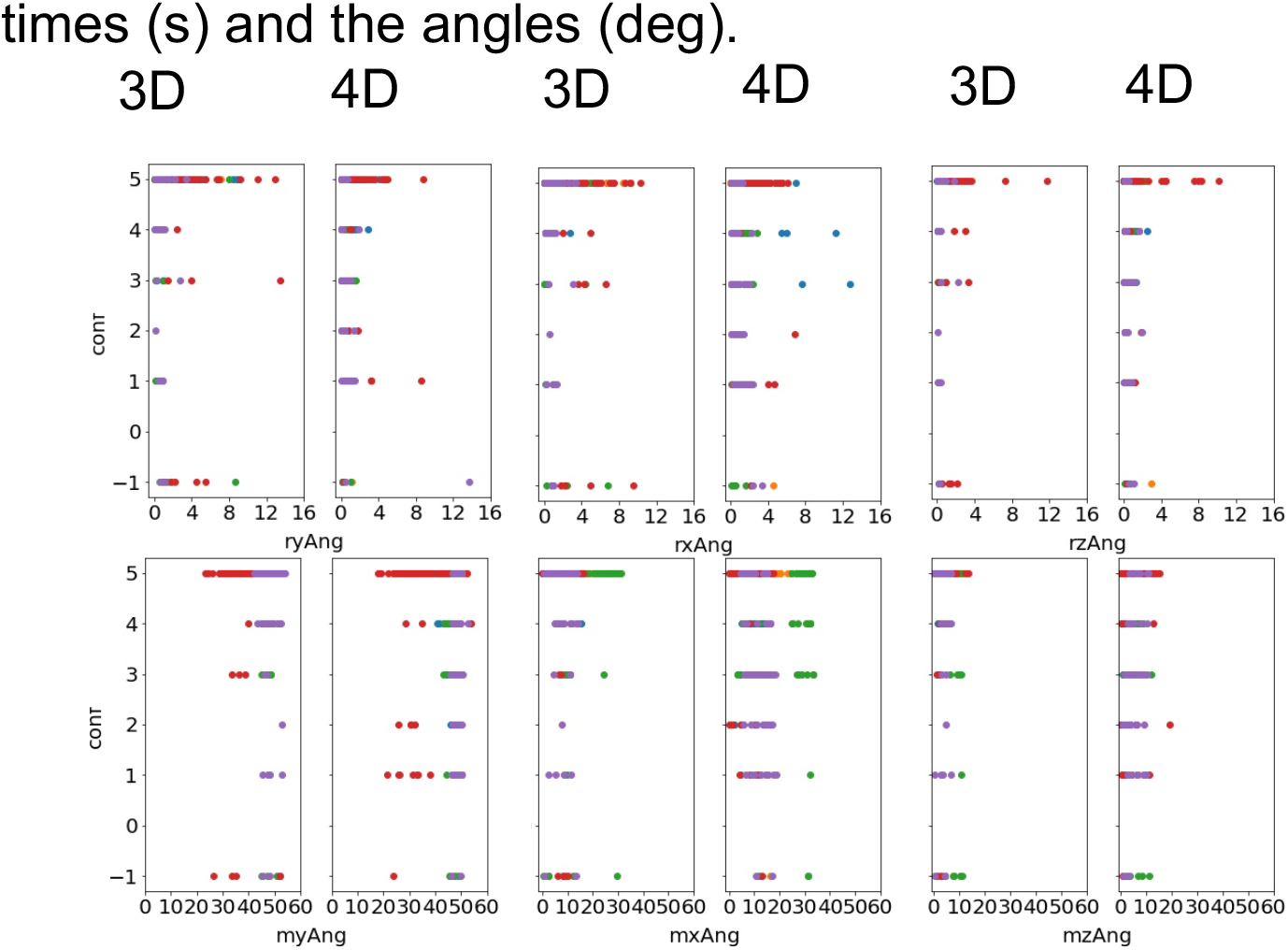
The correlation between the confidence levels (1 to 5) and the angles (deg). The confidence levels of “−1” were from the missed trials.

**Table 3:** Correlations between the Headset Viewpoint and the Performance: Pearson Correlation, p-value.

|  |  | 3D |  |  | 4D |  |  |
| --- | --- | --- | --- | --- | --- | --- | --- |
|  |  | Accuracy (%) | Reaction Time (ms) | Confidence | Accuracy (%) | Reaction Time (ms) | Confidence |
| Range | x | 0.14, <0.01 | 0.18, <0.01 | -0.09, 0.05 | 0.06, 0.2 | 0.19, <0.01 | -0.1, 0.04 |
|  | y | 0.1, 0.03 | 0.1, 0.03 | -0.1, 0.03 | 0.04, 0.41 | 0.1, 0.01 | -0.11, 0.02 |
|  | z | 0.09,<br>0.05 | 0.04,<br>0.37 | 0.01, 0.86 | 0.04,<br>0.37 | 0.02,<br>0.62 | 0.03, 0.6 |
| Median | x | 0.15,<br><0.01 | -0.13,<br><0.01 | 0.07, 0.14 | -0.06,<br>0.26 | 0.2,<br><0.01 | -0.11, 0.03 |
|  | y | -0.29,<br><0.01 | 0.02,<br>0.72 | -0.07, 0.17 | -0.18,<br><0.01 | 0.02,<br>0.74 | -0.09, 0.05 |
|  | z | 0.06, 0.2 | 0.06,<br>0.22 | -0.05, 0.29 | -0.13,<br><0.01 | -0.01,<br>0.84 | -0.08, 0.11 |

As for the median of the angles, we found subjects tended to have their own preferred viewpoints. These viewpoints varied along the x-axis across subjects and spread narrowly along the z-axis within 15 deg, which can be resulted from different heights across subjects and people’s usual horizontal head statics. Interestingly, these angles were generally around 45 deg along the y-axis, which implied a common optimal observation viewpoint selected by the subjects, in spite that there was no strong correlation between the viewpoint and the performance. In fact, the active headset remarkably improved the performance compared to the fixed headset condition and strong ceiling effects were observed. This ceiling performance could weaken its correlation to the headset movement.

## 3. Discussion

Our results show that, given 3D spatial cues, subjects can learn 4D object motion and respond as fast and accurately as for 3D stimulus. This is shown by no significant difference in accuracy and reaction time between 3D and 4D cubes.

With the headset fixed and objects’ vertexes displaced along the x-axis, accuracies in both 3D and 4D recognition are close to perfect and therefore a ceiling effect might exist and reduce the difference resulting by the dimensionality. However, by changing the displacement axis to increase the difficulty, all dependent variables change equally between 3D and 4D conditions, showing a similar dependence on 3D information in both 3D and 4D object motion recognition tasks. Additional support comes from the finding of a compensatory effect on the performance from active exploration or variable viewpoints in both 3D and 4D conditions. With the active headset, subjects’ performance in recognizing objects displaced along difficult axes became as well as the easy axis. Surprisingly, this compensatory effect can be triggered with very slight head movements and viewpoint changes, even in 4D tasks. These findings suggest that subjects can generate 4D phenomenal experiences through 3D displays.

Despite the fact that the x-axis and y-axis shared the same front viewpoint towards the stimuli, subjects showed superiority in performance along the x-axis in the fixed headset condition. This might be due to the fact that our stimuli were shown according to a vertical arrangement, in which one object was shown on the top whereas another one was shown at the bottom. Another possible reason is that displacement along the x-axis causes the vertical orientation change, which has been suggested to convey more binocular disparity information for the stereopsis system [Yazdanbakhsh and Watanabe, 2004].

In addition to increasing the difficulty of tasks at a spatial-dimensional level, an object structural-level factor, irregularity, was also studied. However, it was found to have no effect on the subjects’ performance in these tasks. This implies that subjects’ motion processing in both 3D and 4D tasks was not from shape analysis but spatial speculations. This finding shows that subjects were not recognizing 4D cubes using pure 3D cues without hyperdimensional constructions, since 3D cues are shape-level information for the 4D objects.

In summary, our results add to the corpus of findings showing humans’ ability to perceive higher-dimensional visual stimuli in a broad sense. In terms of phenomenology, one can compare the 4D percepts induced thus far to those of 3D percepts emerging from 2D stimuli: For example, by using perspective cues, we can “perceive” 3D structure from 2D drawings. That percept is not as rich as one that we would get in a VR set generating 3D percepts using disparity and motion cues. What remains to be seen is whether one can generate 4D percepts similar to 3D percepts in a VR by using extensive training.

## Appendix I

The stimuli used in this study included the 3D cubes and 4D tesseracts in different levels of irregularity that were constructed by vertexes and edges connecting them. In the following description, we use the vector, *c_k_*, to indicate the coordinate of a cube or a tesseract’s vertex *k* (cube: *k* = {1,2,…,8}; tesseract: *k* = {1,2,…,16}) and thus define the indexes 0, 1, and 2 that correspond to the *x, y*, and *z* axes respectively. For the tesseract, an additional axis, w, was added, which corresponds to the index 3 in all the vectors and matrixes as follows. Thus, all the vectors in 3D space have three elements, whereas they have four elements in 4D space.

Here we define two basic matrix transformations used in this section: rotation and shear in Equation 1 and 2 respectively. For an n-dimensional object, these matrixes are *n* × *n*. Let *R_ab_*(*θ*) to indicate the coordinate change due to the positive rotation from axis *b* to axis *a* relative to the origin for *θ*.

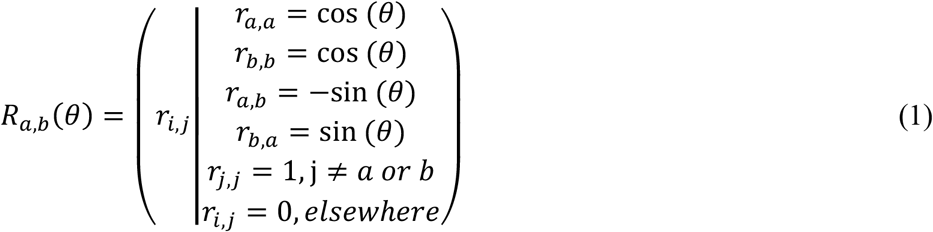

It can be easily verified that *R_x,y_*, *R_y,z_*, and *R_z,x_* respectively indicate to positive rotation around *z, x*, and *y* axes in the 3D space. Extended to 4D space, *R_x,w_*, *R_y,w_*, and *R_z,w_* respectively correspond to the positive rotation around y-z, x-z, and x-y planes.

Also, axial displacement via shearing was used in the non-rigid motion. For example, as illustrated in Figure 3 and 4, by shearing y-axis to x-axis, left-right (x-axis) displacement is created. Similarly, in-out (z-axis) displacement and up-down (y-axis) displacement were created by shearing y-axis to z-axis and shearing z-axis to y-axis respectively. The transformation matrix shearing axis *b* to axis *a* with a degree of *σ* can be determined by Equation 2.

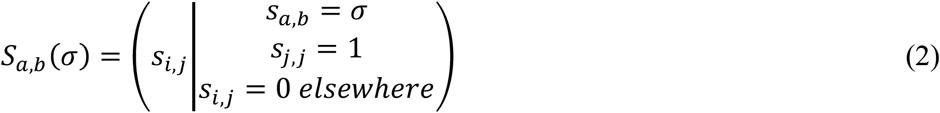

### I.1 3D Rigid Motion

The rigid motion of the 3D cubes was practically realized by adding the random vertex movements and the rotation along *x* and *y* axes. The random movements for the vertex *k, u_k_*, was a vector whose three elements were independently generated from a uniform distribution U(−0.1,0.1).

Therefore, during rigid motion of a 3D cube, the vertex *k*’s movement with respect to time, 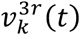, can be determined by the Equation 3.

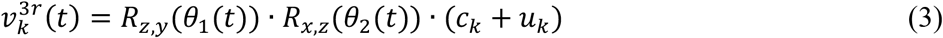

where θ_1_(*t*) and θ_2_(*t*) are smooth random time series generated with two independent Gaussian processes that are subject to *N*(0,1).

### 1.2 3D Non-Rigid Motion

The non-rigidity of the stimuli’s motion was created by firstly deforming the shape of the object, which was achieved by adding random positional shifts, *f_k_*, to the vertexes of stimuli, as indicated by Equation 4. This random shift was synchronized with the rotation around the y-axis to avoid creating obvious discrimination cues.

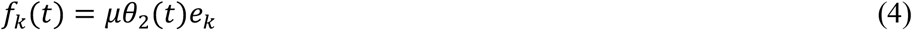

where *μ* is a constant indicating the degree of synchronization, and *e_k_* is a unit vector whose direction is uniformly distributed across the entire 3D space.

After the deformation, an axial displacement was added via shearing one axis to another. Importantly, the degree of shearing in the stimuli was also synchronized with the rotation around the y-axis. Therefore, the vertex *k*’s movement with respect to time during non-rigid motion that shears axis *b* to axis *a* can be represented by Equation 5.

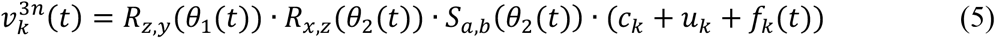

### I.3 4D Motion

Similar to the 3D rigid motion, the 4D tesseracts’ rigid motion were realized by adding the random vertex movements and the rotation along y-z, x-z, and x-y planes. Therefore, the vertex *k*’s movement with respect to time during rigid motion, 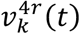, can be determined by the Equation 6.

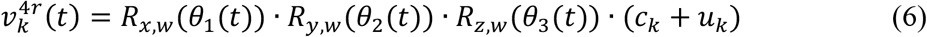

Moreover, the vertex *k*’s movement with respect to time during non-rigid motion that shears axis *b* to axis *a* can be represented by Equation 7.

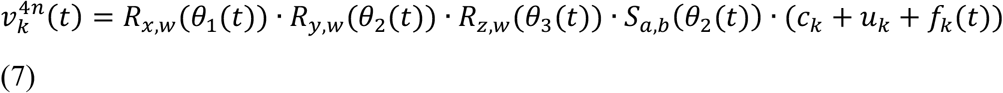

